# Cldn18 evolution reveals pleiotropic constraints in vertebrate stomach loss

**DOI:** 10.64898/2026.09.30.755615

**Authors:** André M. Machado, André Santos, João Moutinho, João Teixeira, Raquel Ruivo, Elsa Froufe, Patrícia G. Ferreira, Mónica Lopes-Marques, Jonathan M. Wilson, L. Filipe C. Castro

## Abstract

Understanding the origins of anatomical innovations is central to elucidate the evolution of phenotypic diversity, as novel body plans often underpin the colonization of new ecological niches. The stomach, with its acid-secreting glands, evolved in the gnathostome ancestor and has been lost repeatedly. Claudin 18 (CLDN18) is a tight-junction protein that preserves gastric epithelial integrity against extreme H^+^ gradients and, in mammals, also acts in the lung through alternative first-exon isoforms. This dual role suggests that pleiotropy may shape the fate of CLDN18 after stomach loss. Here, we provide a detailed analysis of CLDN18 evolution across jawed vertebrates. CLDN18 originated in the gnathostome ancestor and is expressed in the stomach of elasmobranchs. We found no evidence of the gene in chimaeras or lungfishes, two additional agastric lineages, extending the loss of *cldn18a* previously described in agastric teleosts. Exon 1 architecture indicates a tandem duplication in the Euteleostomi ancestor, whereas stomach- and lung-associated isoforms are partitioned in most tetrapods examined. Monotremes, although agastric, retain only the lung-associated exon 1, consistent with lung function sustaining selection on the *locus*. We show that the origin of the stomach in the gnathostome ancestor coincided with the acquisition of a specific tight-junction barrier against extreme H^+^ gradients, a function secondarily lost together with the stomach in chimaeras, lungfishes and agastric teleosts; except in monotremes, where the lung-associated exon 1 preserves CLDN18. Together, these findings reveal a complex evolutionary history involving gene loss, alternative splicing, tissue-specific regulation, and secondary exon loss, highlighting how pleiotropy and regulatory partitioning shaped the evolution of epithelia in jawed vertebrates.

## Introduction

Deciphering the molecular basis of phenotypes is a central goal in biology (Orgogozo et al., 2015). Phenotypic diversity emerges through multiple evolutionary processes, including gene duplication, gene loss, and alternative splicing. Gene duplication has long been recognized as a major source of innovation, allowing paralogues to acquire new or specialized functions (Conrad and Antonarakis, 2007; Kuzmin et al., 2022). Whole-genome duplications are a classic example of this process and have played a major role in vertebrate evolution (Ohno, 1970; Conant and Wolfe, 2008; Zhu et al., 2026). By contrast, the adaptive importance of gene loss has been less widely appreciated, although it is now clear that gene loss can also drive physiological specialization and phenotypic diversification (Olson, 1999; Albalat and Cañestro, 2016; Sánchez-Serna et al., 2025; Valente et al., 2026). In parallel, alternative splicing expands functional diversity by generating distinct isoforms from a single gene and may help reduce pleiotropic constraints through tissue-specific expression (Bush et al., 2017; Tapial et al., 2017; Marasco and Kornblihtt, 2023; Verta and Jacobs, 2022). Together, these processes provide complementary routes to phenotypic evolution. How they combine to shape complex traits, however, remains less clear. Here, we examine this broad question in the context of vertebrate stomach evolution. The stomach is a specialized region of the digestive tract defined by an acidic environment and peptic digestion, both of which depend on the specific expression of the gastric proton pump subunits (ATP4A and ATP4B) and of pepsinogens (Kim and Shivdasani, 2016). I ts developmental program is genetically complex and deeply conserved (Kim and Shivdasani, 2016; Li et al., 2018; Gonçalves et al., 2019). From an evolutionary perspective, the acid-peptic stomach likely originated in the gnathostome ancestor (Koelz, 1992); yet it has been repeatedly lost in several vertebrate lineages, together with its defining gastric glands and associated molecular machinery (Figure 1 A) (Ordóñez et al., 2008; Castro et al., 2014; Gonçalves et al., 2016; Esfandiyari et al., 2022; German, 2024; Kato et al., 2024; Ferreira et al., 2025). In the present work, we examine Claudin 18 (*CLDN18*), a tight-junction protein essential for gastric epithelial integrity (Figure 1B). CLDN18 helps maintain the paracellular barrier of the stomach epithelium, preventing H^+^ leakage under highly acidic conditions and a massive gradient (Figure 1B) (Hayashi et al., 2012). In mice, its loss disrupts tight junction structure and causes atrophic gastritis, underscoring its functional importance (Hayashi et al., 2012). However, in mammals, *CLDN18* has an additional layer of functional complexity. Alternative use of exon 1 generates tissue-specific isoforms expressed in the stomach and lung alveolar epithelium, respectively (Figure 1C) (Niimi et al., 2001; Türeci et al., 2011). Consistent with a pulmonary role, *CLDN18* deficiency in mice also disrupts alveolar barrier function and impairs alveologenesis (LaFemina et al., 2014). This dual gastric–lung expression profile suggests that pleiotropy may have influenced the evolutionary fate of the gene, as highlighted by the contrasting patterns described in agastric vertebrates: the teleost-specific paralogue *cldn18a* is consistently absent in independent agastric teleost lineages, whereas in agastric monotremes *CLDN18* is retained (German, 2024; Kato et al., 2024).

**Figure 1.**
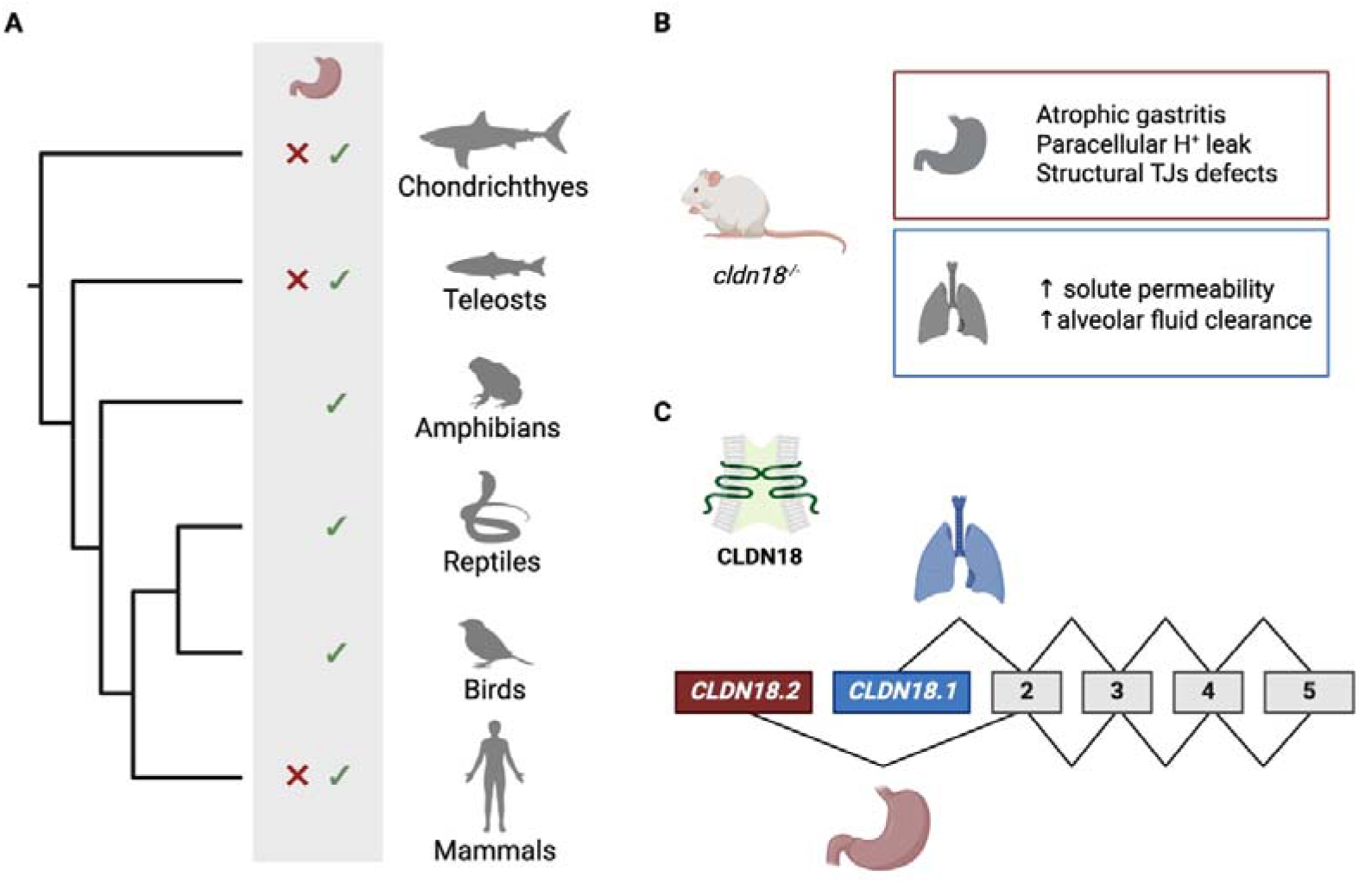
Functional and evolutionary framework of *CLDN18* in gnathostomes. **(A)** Phylogenetic distribution of the stomach across jawed vertebrates. Crosses (X) indicate secondary loss of gastric epithelia, whereas check marks indicate its presence. **(B)** Functional studies in mice show that loss of **Cldn18** (Cldn18^-/-^) disrupts tight junction integrity in the stomach, causing paracellular H leak and atrophic gastritis, whereas in the lung it increases solute permeability and impairs alveolar fluid homeostasis. **(C)** Schematic representation of the CLDN18 gene structure and its major exon 1 splice variants. In mammals, alternative use of exon 1 generates two main isoforms: a gastric-associated form (**CLDN18.2**, red) and a lung-associated form (**CLDN18.1**, blue).

This evolutionary setting raises a central question: how have the origin and diversification of *CLDN18*, particularly its alternative first-exon architecture, shaped its evolutionary fate following recurrent loss of the gastric phenotype? To address this question, this study investigates (1) when *CLDN18* originated during vertebrate evolution; (2) when alternative exon 1 usage emerged; and (3) how gastric- and lung-associated *CLDN18* isoforms are distributed across gastric and agastric gnathostome lineages. Our findings reveal a complex pattern of gene and exon loss consistent with gastric phenotypic variation across extant vertebrates.

## Material and Methods

### Sequence collection and phylogenetic analysis

Human (*Homo sapiens*; NP_001002026.1) CLDN18 amino acid sequence was used as query in BLASTP searches against the NCBI non-redundant protein database. Retrieved sequences were manually inspected for completeness and redundancy. Partial sequences were excluded, except for cldn18 sequences from the chondrichthyan species *Amblyraja radiata, Leucoraja erinacea, and Scyliorhinus canicula*, which were manually curated prior to analysis (Supplementary Table 1). The *Hydrolagus affinis* sequence was manually annotated from an available genome GCA_012026655.1 (Fonseca et al., 2020) using the elephant shark sequence as a reference. A total of 70 amino acid sequences (Supplementary Table 1), representing all major vertebrate lineages and including sequences annotated as CLDN15 as outgroup, were aligned using MAFFT (Katoh et al., 2013). The final alignment comprised 331 positions. Phylogenetic reconstruction was performed using the W-I Q-TREE (Trifinopoulos et al., 2016; Nguyen et al., 2015). The best-fitting amino acid substitution model was determined by the built-in ModelFinder (Kalyaanamoorthy et al., 2017) resulting in JTT+I +G4, and branch support was estimated using the aBayes method (Anisimova et al., 2011). The resulting phylogenetic tree was visualised and annotated in iTOL (Letunic et al., 2021).

### Comparative synteny analysis

To assess conservation of the *CLDN18* genomic region across vertebrates, comparative synteny analyses were performed using representative genome assemblies from major gnathostome lineages. Genomic regions containing *CLDN18*, or the corresponding flanking genes when *CLDN18* was not detected, were identified using a combination of annotation-based gene searches and BLAST similarity searches against NCBI and available genome browsers. Neighbouring genes were recorded and their relative order compared across species. Synteny maps were manually assembled using conserved flanking genes as positional anchors. This approach allowed orthology assessment of candidate *CLDN18* sequences and inference of lineage-specific gene loss when the gene was absent from an otherwise conserved genomic interval. In cases of ambiguous annotation, *locus* structure and neighboring gene content were further inspected to distinguish true orthologues from annotation errors or related claudin family members.

### Identification of exon 1 variants

To investigate the evolutionary distribution of *CLDN18* exon 1 splice forms, genomic sequences and transcript annotations were examined across representative vertebrate species. Exon-intron structures were obtained from annotated gene models whenever available and refined by manual inspection. Candidate exon 1 variants were identified based on their alternative position upstream of shared downstream coding exons and confirmed through sequence comparison with known mammalian gastric- and lung-associated first exons. In species lacking complete transcript annotation, exon 1 candidates were inferred from genomic structure and sequence similarity to orthologous regions in closely related taxa (Supplementary Data 1). Exon 1 alignments were performed in MAFFT (Katoh and Standley, 2013) and graphically represented with ESpript 3.2 (Robert and Gouet, 2014).

### Expression analysis

To infer the tissue-specific expression patterns of *CLDN18* and its alternative exon 1 variants, we surveyed publicly available transcriptomic resources and annotated transcript datasets for representative species (see Supplementary Tables 2, 3 and 4). Genome assembly and annotation files (.fa and .gff) were obtained for 17 species, together with RNA-Seq datasets from up to 12 tissues where available (lung, stomach, brain, heart, kidney, liver, spleen, pancreas, intestine, muscle, testis and ovary; Supplementary Table 2). Expression in lung and stomach is reported for all species (Figure 5; Supplementary Table 3) and in the remaining tissues for ten species (Supplementary Table 4). Relative gene expression was quantified using a standardised pipeline: genomic annotation files were first processed with AGAT (v.1.0.1 ; agat_convert_sp_gxf2gxf.pl; Dainat, 2024), a genome index was built using hisat2-build from HISAT2 software (v.2.2.1) and RNA-Seq reads were aligned with HISAT2 software using the default parameters (Kim et al., 2019). Read abundances were subsequently estimated with StringTie (v.2.1.2; Pertea et al., 2015) using the parameters: -e -B -p 30 -G. Gene-level expression values, reported as transcripts per million (TPM), were extracted from the .gene_abund.tsv output files. For species with evidence of multiple exon 1 variants, tissue distribution was compared to determine whether individual isoforms exhibited restricted or preferential expression, based on the .gtf files generated by StringTie for each sample. Expression assignments were considered conservative and were only reported if supported by transcript annotations or unambiguous RNA-Seq evidence.

## Results and Discussion

A key functional requirement of the stomach epithelium is the strong barrier formed between adjacent cells, which prevents the leakage of H^+^. Tight junctions are central to this paracellular barrier. CLDN18 has a critical role in maintaining this function, as *Cldn18*^*-/-*^mice develop atrophic gastritis due to increased paracellular H^+^ leak (Hayashi et al., 2012). In teleosts, the recurrent loss of *cldn18* is strongly associated with the evolution of the agastric phenotype (Kato et al ., 2024). In contrast, monotremes, which are also agastric, retain an apparently functional copy of the gene (Ordóñez et al., 2008; Dann and Grützner, 2025; Kato et al., 2024). Here we expand the analysis and show that *cldn18* emerged in the gnathostome ancestor and that recurrent loss of the gastric phenotype was associated with either complete gene loss or selective loss of the gastric-associated alternative exon, as observed in monotremes.

### Cldn18 is lost in Chimaeriformes and Dipnoi

We began by surveying the broad diversity of gnathostome lineages spanning both gastric and agastric phenotypes. To reconstruct the phylogenetic distribution and composition of the *CLDN18* gene family, we examined genome assemblies from key representatives across the clade, including cartilaginous fishes (sharks, skates, and chimaeras), Teleostei, Dipnoi (lungfishes), amphibians, reptiles, birds, and mammals (Figure 2; Supplementary Table 1).

**Figure 2.**
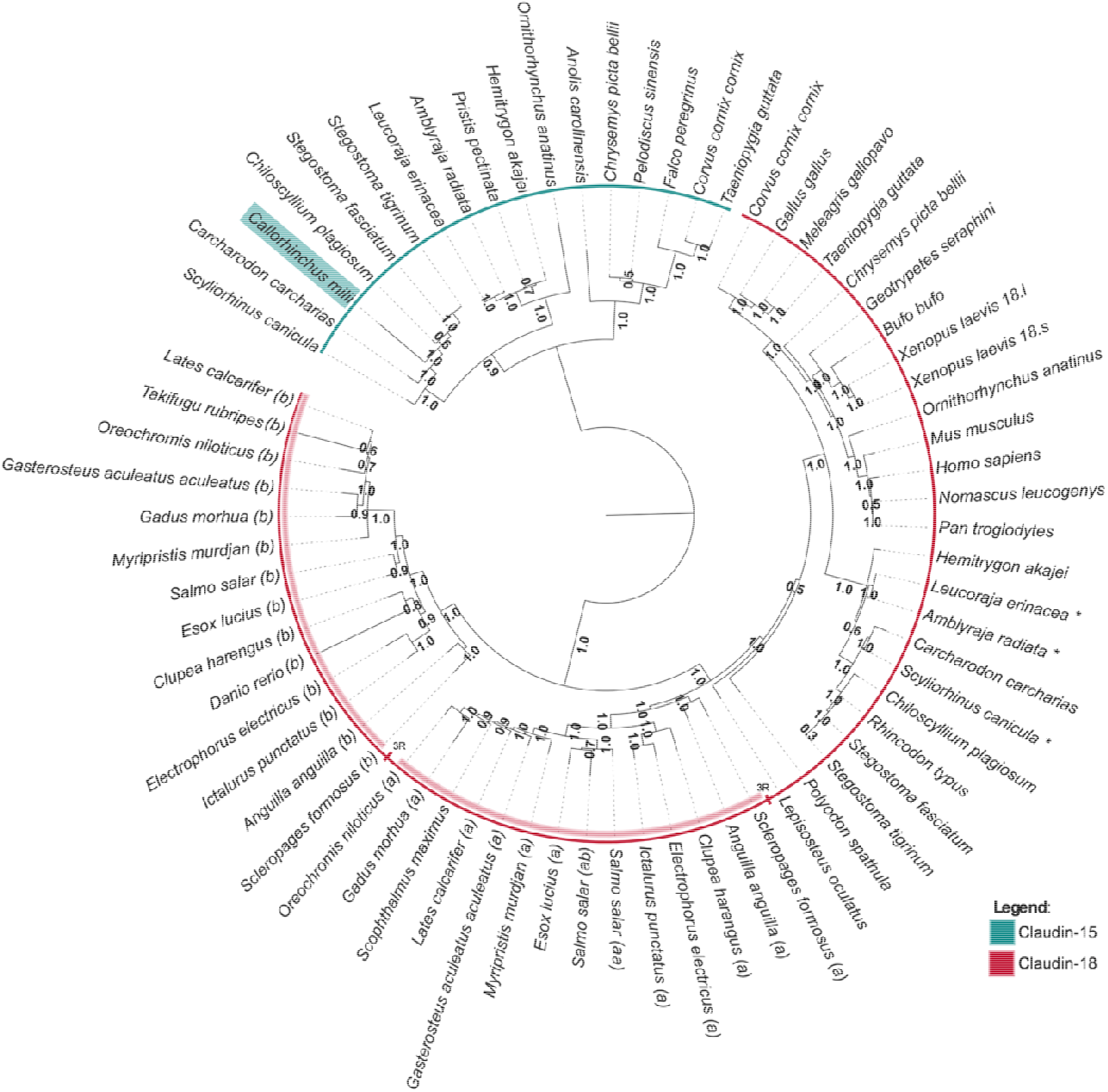
Likelihood phylogenetic analysis of Claudin amino acid sequences, node values indicate aBayes branch support. The C. milii sequence identified as cldn18 in NCBI is highlighted in cyan. Sequences manually curated are indicated with *. Accession numbers of al l sequences are available in Supplementary Table 1. Wider red band indicates cldn18 3R teleost paralogues.

As expected, annotated *cldn18* genes were recovered from gastric sharks and skates. Unexpectedly, however, the elephant shark, an agastric chimaera species (Castro et al., 2014), also carries a *cldn18* annotation (NP_001279939.1; XP_007885328.1) in the current genome assembly and annotation (GCF_018977255.1). To test the orthology of these annotated sequences, phylogenetic analysis was conducted including the putative *cldn18* sequences and the closely related *cldn15* family. The resulting tree resolved two distinct groups: one containing bona fide *cldn18* orthologues and a second containing *cldn15* sequences (Figure 2). The *cldn18* clade included sequences from gastric lineages but notably excluded the annotated elephant shark *‘cldn18’* sequence. Instead, the chimaera sequence grouped robustly within the *cldn15* clade (Figure 2), indicating that the elephant shark *cldn18* is misannotated and is instead a *cldn15* gene.

To further support the orthology analysis, we next examined the *locus* synteny in cartilaginous fishes (Figure 3). Despite overall conservation of the surrounding genomic region, *cldn18* was absent from the elephant shark locus, whereas it was clearly present in elasmobranchs (Figure 3). This analysis was extended to a second chimaera, *Hydrolagus affinis*, using a low-coverage genome assembly (Fonseca et al., 2020), and again no evidence of *cldn18* was found (Figure 3). Together, these results indicate that, as expected and similarly to agastric teleosts, chimaeriforms lost *cldn18* in the common ancestor of the two species examined, consistent with the absence of gastric glands in the lineage. The data also show that the sequence currently annotated as *‘cldn18’* in chimaeriforms is in fact an orthologue of *cldn15*, indicating that the existing annotation is erroneous.

**Figure 3.**
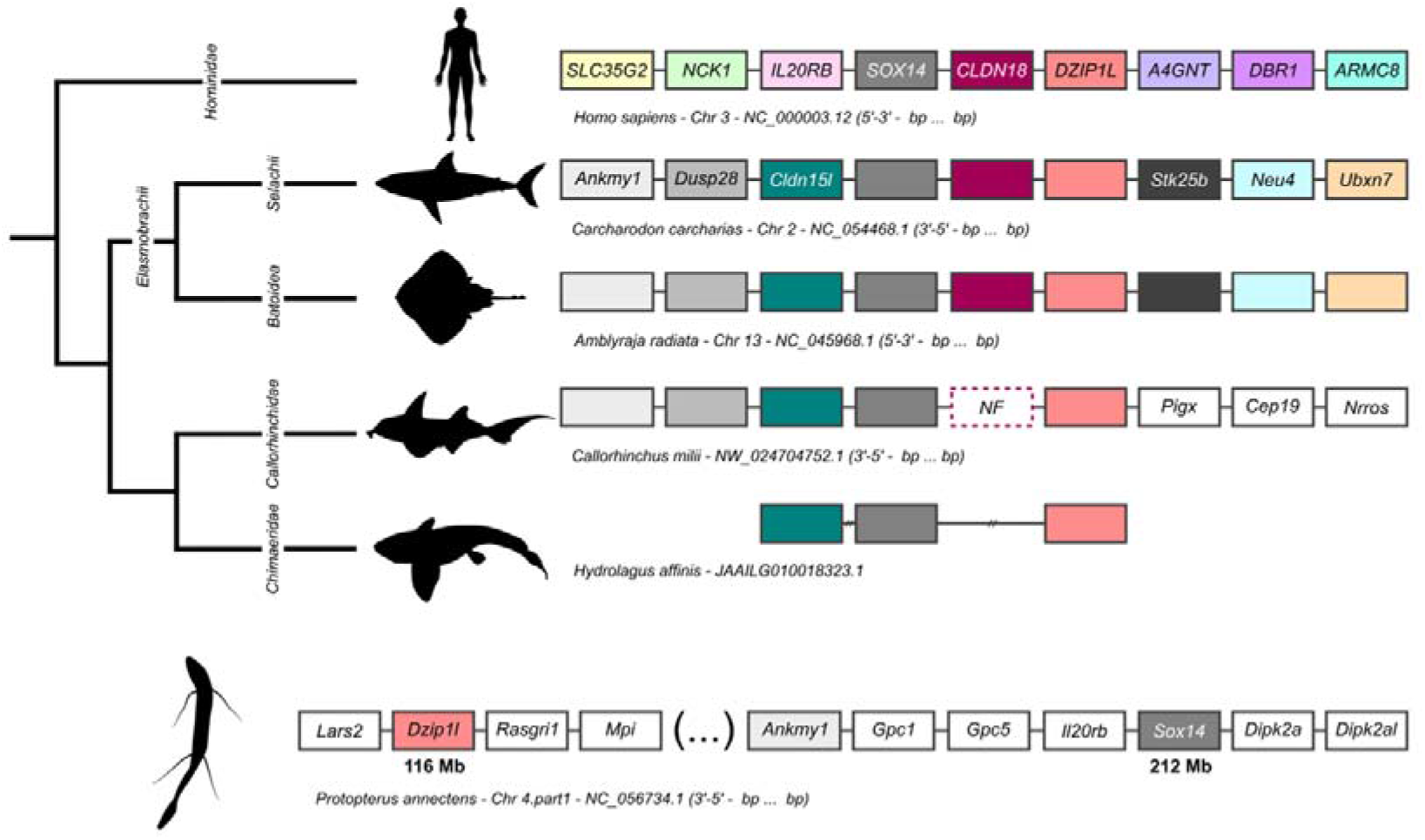
Comparative synteny of the cldn18 locus in gnathostomes. Orthologous genes are shown as colored boxes, with neighbouring genes in grey, to illustrate conservation of local gene order and orientation. Coordinates refer to the genomic interval displayed for each species. This synteny framework was used to assess locus conservation, confirm orthology, and identify lineage-specific absence of cldn18. “NF” denotes that no gene was found at the expected syntenic position.

We next examined a second agastric lineage in which gene loss has also been proposed, the Dipnoi (Castro et al., 2014). Although syntenic conservation around the *cldn18* locus was not preserved, we found no evidence for the presence of *cldn18* in lungfishes (Figure 3). Together, these results show that two additional gnathostome lineages lacking gastric glands, namely chimaeriforms and lungfishes, appear to have lost *cldn18*, extending the pattern previously described in agastric teleosts (Kato et al., 2024).

### The cldn18 exon1 duplication predates Euteleostomi and exon loss correlates with the absence of stomach in monotremes

Our findings suggest that *cldn18* loss correlates with the absence of gastric glands in chimaeras and lungfishes, extending the pattern previously described in agastric teleosts (Kato et al., 2024). Yet monotremes, despite their well-established loss of stomach, clearly retain an orthologue of *CLDN18* (Ordóñez et al., 2008; Kato et al., 2024). This contrast raises the possibility that the evolutionary fate of *CLDN18* differs according to potential functional partitioning. In mammals, *CLDN18* is known to produce alternative exon 1 splice forms with stomach- and lung-restricted expression profiles, a pattern subsequently confirmed across multiple mammalian orthologs (Niimi et al., 2001 ; Türeci et al., 2011). However, the phylogenetic distribution of *CLDN18* splicing, and expression patterns across gnathostomes, remain unknown. We therefore investigated the occurrence of exon 1 alternative splicing of *CLDN18* across major gnathostome lineages (Figure 4).

**Figure 4.**
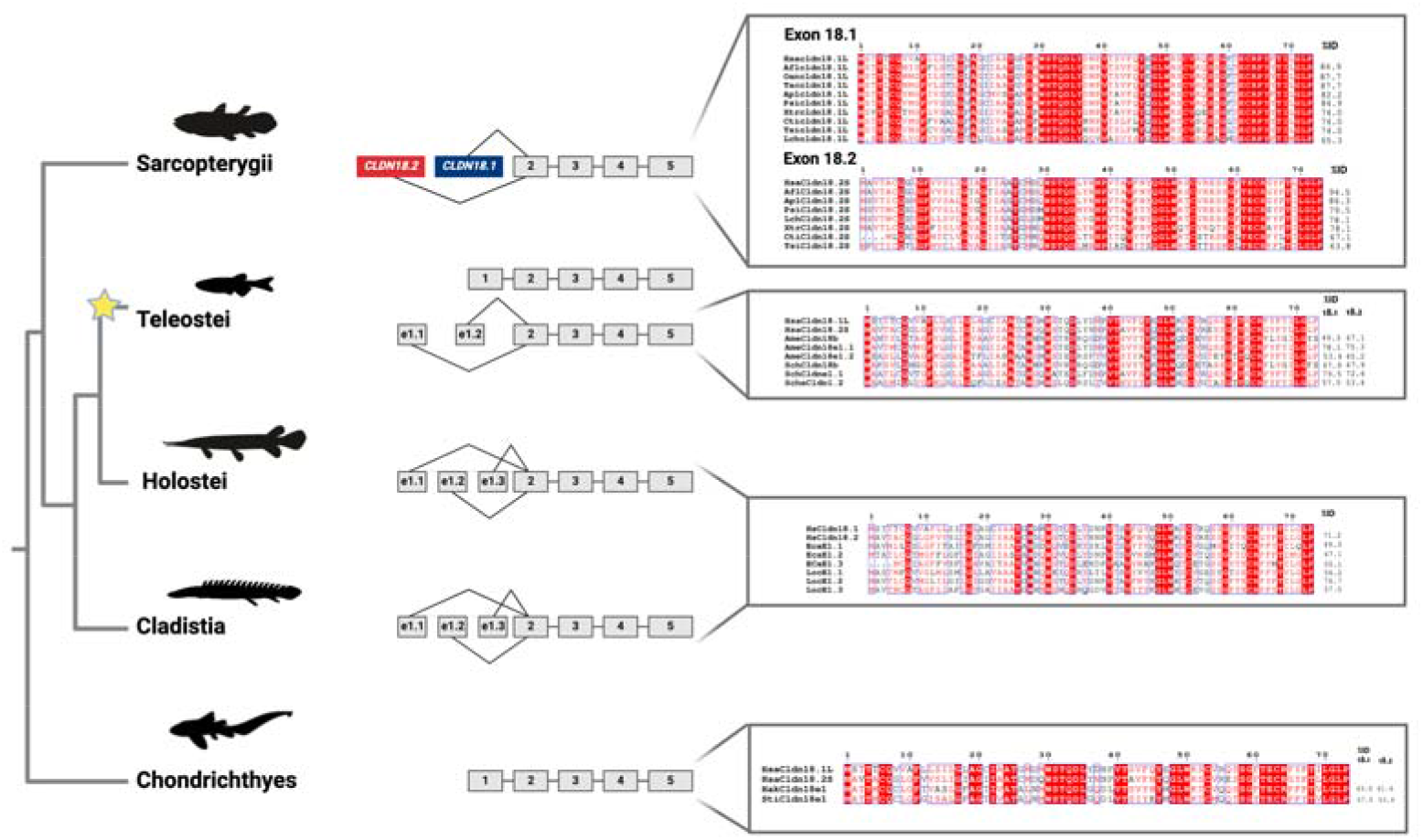
Phylogenetic distribution and pairwise sequence identity of CLDN18 exon 1 splice variants across gnathostome lineages. Gene models (centre) and exon 1 amino-acid alignments (right) are shown for each lineage of the cladogram (left); numbered boxes are coding exons, and the star marks the teleost-specific genome duplication (3R). Sarcopterygii: two alternative first exons, 18.2 (stomach-associated, red) and 18.1 (lung-associated, blue), spliced to shared exons 2–5, with alignments of the lung-associated (Exon 18.1, top) and stomach-associated (Exon 18.2, bottom) sequences from representative sarcopterygian taxa. Teleostei: *cldn18b* (upper gene model, single exon 1) and *cldn18a* (lower gene model, alternative first exons e1.1 and e1.2), the two 3R-derived paralogues, with alignments of exon 1 sequences from representative gastric teleosts. Holostei and Cladistia: three alternative first exons (e1.1, e1 .2, e1.3) sharing exons 2–5, with alignments of the exon 1 sequences of spotted gar and reed fish. Chondrichthyes: single exon 1, with the alignment of the sequences from cartilaginous fishes (Hak, Sti). Pairwise % identity to the human reference exons (HsaCldn18.1L and HsaCldn18.2S) is given to the right of each sequence, and conserved residues are shaded in red. Species abbreviations and accession numbers are given in Supplementary Table 1. *Hsa=Homo sapiens, Oan=Ornithorhynchus anatinus, Tac=Tachyglossus aculeatus, Xtr=Xenopus tropicalis, Lch=Latimeria chalumnae, Eca=Erpetoichthys calabaricus, Loc=Lepisosteus oculatus. Hak=Hemitrygon akajei, Sti=Stegostoma tigrinum*.

In Chondrichthyes, *cldn18* harbours a single exon 1 with expression restricted to the stomach (no more than 0.2 TPM in any other tissue; Figures 4 and 5; Supplementary Tables 3 and 4), representing most likely the ancestral state. Pairwise identity between the two chondrichthyan sequences is 53–63% to either human isoform, with no bias toward human lung-or stomach-associated exons, consistent with a pre-partitioning ancestral state. In Sarcopterygii, *cldn18* displays two alternative first exons across all examined taxa, including the coelacanth (for which transcriptomic data is unavailable) (Figure 4). The distal exon (18.2) is the stomach-associated form, and the proximal exon (18.1) is the lung-associated form (Figures 4 and 5), with within-clade identities to the human references averaging ∼73% for 18.2 (range 61–94%) and ∼78% for 18.1 (range 62–99%). The only exception is the snake lineage, which retains both exons but lacks lung expression in our dataset (Figure 5; Supplementary Table 3), possibly reflecting lineage-specific reorganisation of the pulmonary epithelium within a remarkably diverse group (Hill et al., 2025). Outside the lung and stomach, expression was low or undetectable in all species (no more than 2.4 TPM), except in the intestine of mouse and Xenopus and the kidney and testis of Xenopus (Supplementary Table 4). In monotremes, only a single exon 1 is detected, with high sequence similarity to the mammalian lung-associated form (87.7% to HsaCldn18.1L for both platypus and echidna), indicating secondary loss of the gastric exon while preserving the lung-associated isoform.

**Figure 5.**
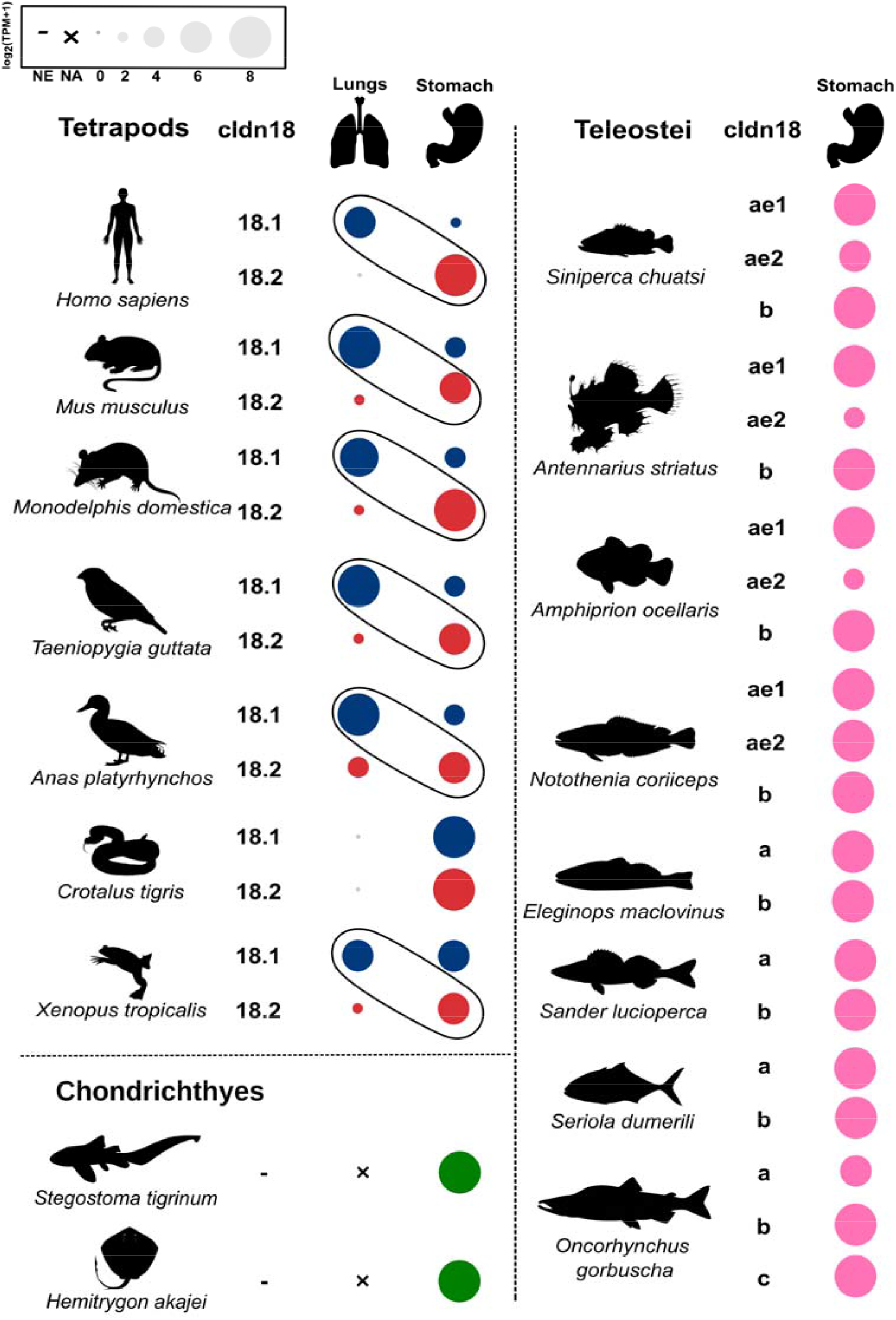
Relative expression of cldn18 first-exon isoforms in the lung and stomach of 17 vertebrate species: seven tetrapods, eight teleosts and two chondrichthyans. Circle area is proportional to log_2_(TPM+1). In tetrapods, 18.1 (blue) and 18.2 (red) denote the lung- and stomach-associated exon 1 isoforms, respectively, and ovals link the two isoforms where their expression is partitioned between lung and stomach. In teleosts (pink), a and b denote the paralogues cldn18a and cldn18b, and ae1 and ae2 the two alternative first exons of cldn18a (e1.1 and e1.2 in Figure 4), where annotated; three paralogues (a, b, c) are shown for Oncorhynchus gorbuscha. In chondrichthyans (green), which have a single exon 1, one value is shown per species. NE, not examined; NA, not applicable (organ absent). Expression values, including those of the remaining tissues analysed, are given in Supplementary Tables 3 and 4.

In early-branching Actinopterygii, Cladistia (reed fish) and Holostei (spotted gar), *cldn18* harbours three alternative first exons arranged in tandem (e1 .1, e1.2, e1.3; Figure 4). Within-clade identities among these six variants are low (∼40–60%), indicating ancient diversification; only one variant (Loce1.2 in gar) approaches the sarcopterygian identity range against the human references (76.7% to HsaCldn18.1L, 74.0% to HsaCldn18.2S), while the remaining five span 47–67% and cannot be assigned to a specific tetrapod counterpart on sequence alone. Expression data for these lineages are presently not available. Importantly, in Teleostei, the two 3R-derived paralogues differ in architecture: *cldn18a* (the gastric form; Kato et al., 2024) retains two alternative first exons (e1.1 and e1.2), while *cldn18b* presents a single exon 1 (Figure 4). Pairwise identity to the human references does not permit unambiguous ancestry assignment: *cldn18a* e1.1 sits within the sarcopterygian range (72–80% to both HsaCldn18.1L and HsaCldn18.2S), e1.2 is markedly more divergent (45–58%), and *cldn18b* is essentially equidistant from both references (48–51% to either; ≤2% differential). In the gastric mandarin fish (*Siniperca chuatsi*), both alternative exon 1 variants of *cldn18a* are expressed in the stomach and are undetectable in the six other tissues available (Supplementary Tables 3 and 4), and *cldn18b* is also expressed in the stomach (Supplementary Table 3). In the seven further teleosts analysed, for which only stomach data are available (Figure 5; Supplementary Table 3), all paralogues and alternative first exons are also expressed in the stomach, and the salmonid *Oncorhynchus gorbuscha* carries a third paralogue (c).

Taken together, these patterns are most parsimoniously explained by a single tandem duplication of *cldn18* exon 1 in the Euteleostomi ancestor, generating a minimal set of a two-copy exon 1 array that was subsequently modified by lineage-specific duplication and loss: yielding three exons in basal Actinopterygii; two in Sarcopterygii and in teleost *cldn18a;* and, one in teleost *cldn18b*. Under this model, tissue-restricted partitioning of the alternative exons into stomach and lung expression emerged in tetrapods as a regulatory innovation layered onto an already-established multi-exon framework, with the distal-to-exon-2 position consistently corresponding to the ancestral stomach-associated function.

### Alternative splicing, gene loss, and organ evolution in gnathostomes

Our results place *CLDN18* at the intersection of several evolutionary processes that are often considered separately but rarely resolved together within a single *locus*: gene duplication, lineage-specific gene loss, alternative splicing, and secondary exon loss. Accordingly, *CLDN18* provides a particularly clear example of how organ evolution can be traced through the history of a gene family. More broadly, our findings support the view that vertebrate organ evolution is shaped not only by the emergence of new genes or new expression domains, but also by selective gene retention, regulatory partitioning, and repeated gene erosion following phenotypic simplification.

A central inference from our data is that *CLDN18* was ancestrally gastric-specific. In cartilaginous fishes, we identified a single exon 1 form with stomach-restricted expression, consistent with a primary role in maintaining the epithelial barrier of the acid-peptic stomach. This interpretation is coherent, as gastric function depends not only on acid secretion and protease activity, but also on the ability of the epithelium to withstand extreme luminal conditions without paracellular leakage. Under this framework, the repeated loss of *Cldn18* in agastric lineages is unlikely to be incidental (Kato et al., 2024). Rather, it suggests that once the gastric gland system was lost, *CLDN18* became dispensable and was repeatedly released from purifying selection. The pattern documented here in chimaeriforms and lungfishes, together with that previously described in agastric teleosts (Kato et al., 2024), is therefore consistent with a broader evolutionary association between stomach loss and erosion of a core gastric epithelial component.

Simultaneously, the evolutionary history of *CLDN18* cannot be reduced to a simple pattern of stomach-associated gene loss. In teleosts, the family was additionally shaped by the teleost-specific whole-genome duplication (Kato et al., 2024). More importantly, in both actinopterygian and sarcopterygian lineages, diversification of exon 1 generated alternative splice forms. Alternative splicing is increasingly recognized not only as a source of transcript diversity, but also as a mechanism that partitions gene function across tissues and organs thereby changing the selective landscape acting on a *locus* (Verta and Jacobs, 2022). In effect, the history of *CLDN18* suggests that exon 1 innovation expanded the functional range of the gene beyond the stomach, diversifying into new epithelial contexts such as the lung, where the *Cldn18. 1* isoform likely contributes to alveolar barrier integrity and protection against pulmonary edema (LaFemina et al., 2014). As observed for the stomach epithelial barrier, the alveolar blood–gas barrier relies strongly on tight junctions to maintain a thin but mechanically robust interface compatible with efficient gas exchange (Leiby et al., 2020). Consistent with this role, *Cldn18* knockout in mice increases alveolar permeability and alters fluid clearance, demonstrating a nonredundant function for CLDN18 in alveolar barrier integrity and homeostasis (Li et al., 2014). The Sarcopterygian pattern is especially informative. Across the taxa examined, two exon 1 variants are associated with stomach and lung expression domains, indicating that tissue-specific isoform partitioning was already established early in tetrapod evolution, as seen in amphibians, birds and mammals, with the snake as the only exception (Figure 5). Within mammals, this partitioning is reinforced by deep conservation of both isoform-coding sequence and isoform-specific *cis*-regulatory elements — including the CREB binding site that activates CLDN18.2 in gastric epithelia and the T/EBP/NKX2.1 binding sites that activate CLDN18.1 in alveolar cells — across all examined mammalian orthologs (Türeci et al., 2011). This regulatory architecture provides a plausible mechanism for the exon-specific evolutionary decoupling observed in monotremes, where stomach-restricted CREB-dependent activation of the gastric exon could be lost without disrupting NKX2.1-dependent activation of the lung exon (Türeci et al., 2011). This suggests that recruitment of *CLDN18* to the lung was not merely an extension of ancestral expression, but a derived step in which exon-specific regulation created a second organ-linked functional axis. Once *CLDN18* acquired an additional role in the lung, complete deletion of the gene would carry a broader functional cost. The locus would no longer evolve solely under gastric selection, but under the combined constraints of two epithelial systems. Thus, our results support a model in which alternative splicing does not simply add expression complexity, but actively restructures pleiotropy.

An informative consequence of this framework is that exon 1 duplication of *cldn18* is temporally decoupled from its tissue-specific allocation. The duplication itself is apparently ancestral to Euteleostomi, but assignment of the resulting alternative exons to stomach- and lung-restricted expression is most likely tetrapod-specific. Coherently, the paired, vascularised, air-breathing lung is itself a tetrapod-stem innovation (Cupello et al., 2022), even though much of the underlying genetic foundation for pulmonary specialisation can be traced back to gnathostome ancestors (Li et al., 2025). In teleosts, where no tetrapod-alveolar type lung exists, the two alternative *cldn18a* first exons partition not into stomach and lung, but into stomach only in the eight teleosts examined (Figure 5; Supplementary Table 3). The same multi-exon-1 framework was therefore available across Euteleostomi but recruited differently, with lung allocation emerging only when the tetrapod-type lung itself emerged. Expression data are not yet available for the alternative first exons of coelacanth, reed fish, or spotted gar; given that coelacanths represent the sister group to Dipnoi and tetrapods and retain a vestigial lung, and that reed fish and gar retain the ancestral three-exon architecture, these lineages would be highly informative for dating the onset of tissue allocation.

A direct consequence is that alternative splicing alone does not protect a gene from erosion when only a single organ system depends on it. In agastric teleost lineages, *cldn18a* is lost wholesale together with both of its alternative first exons (Kato et al., 2024). The remaining ohnolog, *cldn18b*, is retained in some agastric teleosts but shows increased non-synonymous substitution relative to gastric species (Kato et al., 2024), consistent with its expression in the stomach of the mandarin fish (Figure 5; Supplementary Table 3) and with relaxed constraint after stomach loss. The presence of two splice forms does not buffer the gene against loss, because both forms are expressed in the gastric system. Splice diversity *per se* is therefore insufficient to maintain a *locus* across phenotypic transitions; what is required is functional partitioning across *independent* organ systems that are not jointly eroded by the same phenotypic change.

This framework also offers a possible interpretation of an apparent paradox among the Dipnoi. Lungfishes possess functional lungs and have lost the stomach (Cupello et al., 2022; Castro et al., 2014), yet they have lost *cldn18* entirely. Under a simple “*lung function preserves the gene*” model, retention would be expected. One possibility is that lungfish lungs rely on a different claudin complement for epithelial barrier integrity. Alternatively, the recruitment of Cldn18 to a specialized pulmonary function can be interpreted as a tetrapod-stem innovation rather than a pan-Sarcopterygian feature, with stomach loss in Dipnoi then removing the only ancestrally constrained function. The subsequent acquisition of a lung-associated function in the tetrapod lineage would have provided an additional selective constraint, allowing retention of the gene even with the co-occurrence of gastric function loss. The Dipnoi pattern therefore is compatible with the emergence of pulmonary pleiotropy at the tetrapod node, a hypothesis that remains to be tested with lung expression data from lungfish, coelacanth and early-branching ray-finned fishes, and is consistent with the broader observation that the genetic foundation for the lung predates the appearance of paired lungs themselves (Li et al., 2025).

Our findings also shed light on the monotreme paradox. If *CLDN18* were exclusively associated with gastric function, its retention in platypus and echidna would be difficult to reconcile with their well-established agastric phenotype. Instead, our data indicate that monotremes retained only the exon 1 variant most similar to the mammalian lung-associated form, whereas the gastric-associated exon was secondarily lost. This suggests that *CLDN18* was preserved in monotremes not because gastric function persisted, but most likely because pulmonary function, maintained selection on the *locus*, although expression data for the monotreme lung are not yet available. The monotreme pattern therefore provides an example of exon-specific evolutionary decoupling: stomach loss was followed by loss of the gastric exon, but not by loss of the gene as a whole.

Taken together, our results suggest a broader principle for organ evolution in vertebrates. The molecular consequences of organ loss are not uniform across lineages, because they depend on how strongly a given gene is integrated into other functional systems. Where a gene is largely organ-restricted, phenotypic loss may be followed by full gene erosion, as seen here for *cldn18* in chimaeriforms, lungfishes, and agastric teleosts. Where the same *locus* has been repartitioned through alternative splicing and recruited to another essential tissue, organ loss may instead result in partial, exon-specific remodelling. Thus, *CLDN18* does more than track stomach evolution: it shows how pleiotropy, once reorganized through regulatory innovation, can constrain or redirect the genomic consequences of phenotypic simplification. Overall, our findings identify *CLDN18* as a molecular marker of both organ loss and organ coupling, showing how gene erosion, exon-specific retention, and pleiotropy together shaped the evolution of the stomach and lung in jawed vertebrates.

### Declaration of generative AI and AI-assisted technologies in the manuscript preparation process

During the preparation of this work the author(s) used Claude (Sonnet 5.5) (Anthropic) in order to improve the language and readability of the manuscript, check its internal consistency (citations, figure and table references). After using this tool/service, the author(s) reviewed and edited the content as needed and take(s) full responsibility for the content of the published article.

## Author contributions

L.F.C.C. and J.M.W. conceived and supervised the study. A.M.M., A.S., J.M., J.T. and R.R. performed the sequence retrieval, phylogenetic, synteny and exon 1 analyses. A.M.M.,

J.M. and J.T. carried out the expression analyses. E.F. and P.G.F. contributed data, genomic resources and interpretation. M.L.-M. contributed to the evolutionary analyses and interpretation. L.F.C.C. and J.M.W. wrote the original draft. All authors discussed the results, reviewed and edited the manuscript, and approved the final version.

## Supplementary Tables and Data

Supplementary Table 1. Accession numbers and NCBI protein names of the amino acid sequences used in the phylogenetic analysis (Figure 2).

Supplementary Table 2. Genome assemblies and RNA-Seq datasets (SRA run accessions) used to infer relative gene expression in 17 vertebrate species.

Supplementary Table 3. Relative expression (TPM) of CLDN18 and its alternative exon 1 transcripts in the lung and stomach of 17 vertebrate species.

Supplementary Table 4. Relative expression (TPM) of CLDN18 and its alternative exon 1 transcripts in nine further tissues of ten vertebrate species.

