## Supplemental Material 1 for "Cldn18 evolution reveals pleiotropic constraints in vertebrate stomach loss"

**Supplementary material 1 - Manual annotation of partial Chondrichthyes sequences**

Manual annotated sequences, grey highlighting indicates sequence portion missing from NCBI file that was manually annotated in this work.

>XP_032887427.1 Claudin 18 *Amblyraja radiata*

MATTMCQCLGFTVASLGFVGTIVATALNMWSTQDLGQDLVTSVYNYMGLWKTCVQQTSGFTECRPFFTVL
GLPATFQAIRALMIVSIVLSLIGSMVSILALKCIRLGTMEDTAKANLTLTSGILFIVAGVCCVAGVSIYA
NMLVTNFWMSNTSMLGSQGPARQRYTFGSALFVGWVSGGLLLVGGIVMCIACRGMVPEDTRYNSTIYKAP
SRGHTYKSGYKSSTHYETEMAPVKRTYPDEKSAYNRSTNEDTKQGKYDYV

>XP_055501994.1 claudin 18 like *Leucoraja erinacea*

MATTMCQCLGFTVASLGFVGTIVATALNMWSTQDLGQDLVTSVYNYMGLWKTCVQQTSGFTECRPFFTVL
GLPATFQAIRALMIVSIVLSLIGSMVSILALKCIRLGTMEDTAKANLTLTSGILFIVAGVCCVAGVSIYA
NMLVTNFWMSNTSMLGSQGPARQRYTFGSALFVGWVSGGLLLVGGIVMCIACKGMVPEETRYNSTIYKAP
SRGHTYKSGYKSSTHYETEMAPVKRTYPDEKSAYNRSTNEDTKQGKYDYV

>XP_038671664.1 Claudin18-like *Scyliorhinus canicula*

MATTMCQCLGFGISTLGFAGTIVATALNMWSTQDLGRDLVTTVFKYMGLWKTCLQQTSGFTECRPFYTVL
GLPATFQAVRALMIVAIVLSLIGSGVSIMALKCIRVGTMDDAAKANLTLTSGVLFIVAGVCCIAGVSIYG
NMLVTDFWQMNGSILGGQGSVQERYTFGAALFVGWVSGGLLVIGGIVMCIACRGLMPEDTRYNSTIYKVP
SRGAGYKTGYKASTHYDTEMTHVRKTYPDDVESKKSAYNRSTNEDSKRGAPAKYDYV

**Supplementary table 1:** Accession numbers and protein names of amino acid sequences used in the phylogenetic analysis. *C. milii* Sequences annotated as Claudin 18 in Ncbi are highlighted in red.

| # | Accession number | NCBI protein nomenclature | Species |
|---|---|---|---|
| 1 | XP_072896750.1 | CLDN_15-like | *Hemitrygon_akajei* |
| 2 | XP_051873979.1 | CLDN15-like | *Pristis_pectinata* |
| 3 | XP_055501998.1 | CLDN15-like | *Leucoraja_erinacea* |
| 4 | XP_032887419.1 | CLDN15-like | *Amblyraja_radiata* |
| 5 | GCA_012026655.1 | CLDN15-like | *Hydrolagus_affinis* |
| 6 | NP_001279939.1 | CLDN18 | *Callorhinchus_milii* |
| 7 | XP_038673117.1 | CLDN15-like | *Scyliorhinus_canicula* |
| 8 | XP_041036124.1 | CLDN15-like | *Carcharodon_carcharias* |
| 9 | XP_043557722.1 | CLDN15-like | *Chiloscyllium_plagiosum* |
| 10 | XP_048398239.1 | CLDN15-like | *Stegostoma_fasciatum* |
| 11 | XP_048398239.2 | CLDN15-like | *Stegostoma_tigrinum* |
| 12 | XP_043926512.1 | CLDN15-like | *Protopterus annectens* |
| 13 | XP_028931688.1 | CLDN15-like | *Ornithorhynchus_anatinus* |
| 14 | XP_008104700.1 | CLDN15-like | *Anolis_carolinensis* |
| 15 | XP_005282529.1 | CLDN19 | *Chrysemys_picta_bellii* |
| 16 | XP_006125254.1 | Claudin-19-like | *Pelodiscus_sinensis* |
| 17 | XP_005232313.2 | CLDN15-like | *Falco_peregrinus* |
| 18 | XP_019145079.1 | CLDN-19 | *Corvus_cornix_cornix* |
| 19 | XP_041574102.1 | Claudin-19 | *Taeniopygia_guttata* |
| 20 | NP_062789.1 | CLDN18 | *Mus_musculus* |
| 21 | NP_001002026.1 | CLDN18 | *Homo_sapiens* |
| 22 | XP_526318.3 | CLDN18 | *Pan_troglodytes* |
| 23 | XP_003265323.1 | CLDN18 | *Nomascus_leucogenys* |
| 24 | XP_028925772.1 | CLDN18 | *Ornithorhynchus_anatinus* |
| 25 | XP_033815148.1 | CLDN18 | *Geotrypetes_seraphini* |
| 26 | XP_040286930.1 | CLDN18 | *Bufo_bufo* |
| 27 | NP_001083443.1 | claudin_18_L | *Xenopus_laevis* |
| 28 | XP_018120300.1 | CLDN18 | *Xenopus_laevis* |
| 29 | XP_005303714.1 | CLDN18 | *Chrysemys_picta_bellii* |
| 30 | XP_039413104.1 | CLDN18 | *Corvus_cornix_cornix* |
| 31 | XP_040535069.1 | CLDN18 | *Gallus_gallus* |
| 32 | XP_002196241.1 | CLDN18 | *Taeniopygia_guttata* |
| 33 | XP_003209013.1 | CLDN18 | *Meleagris_gallopavo* |
| 34 | XP_072896748.1 | CLDN18-like | *Hemitrygon_akajei* |
| 35 | Manual annotation | CLDN18 | *Amblyraja radiata* |
| 36 | Manual annotation | CLDN18 | *Leucoraja erinacea* |
| 37 | Manual annotation | CLDN18 | *Scyliorhinus canicula* |
| 38 | XP_041029914.1 | CLDN18-like | *Carcharodon_carcharias* |
| 39 | XP_043557721.1 | CLDN18-like | *Chiloscyllium_plagiosum* |
| 40 | XP_048398771.1 | CLDN18-like | *Stegostoma_fasciatum* |
| 41 | XP_048460550.1 | CLDN18 | *Rhincodon_typus* |
| 42 | XP_048398771.1 | CLDN18-like | *Stegostoma_tigrinum* |
| 43 | XP_006637728.1 | CLDN18 | *Lepisosteus_oculatus* |
| 44 | XP_035272052.1 | CLDN18-like | *Anguilla_anguilla* |
| 45 | XP_018593928.1 | CLDN-18-like | *Scleropages_formosus* |
| 46 | XP_031430652.1 | CLDN18 | *Clupea_harengus* |
| 47 | XP_017334649.1 | CLDN18 | *Ictalurus_punctatus* |
| 48 | XP_013979138.1 | CLDN18 | *Salmo_salar* |

| | | | |
|---|---|---|---|
| 49 | XP_019900146.1 | CLDN18 | *Esox_lucius* |
| 50 | XP_014024098.1 | CLDN18-like | *Salmo_salar* |
| 51 | XP_029905880.1 | CLDN18-like | *Myripristis_murdjan* |
| 52 | XP_040039231.1 | CLDN18-like | *Gasterosteus_aculeatus_aculeatus* |
| 53 | XP_018525411.1 | CLDN-18 | *Lates_calcarifer* |
| 54 | XP_035502666.1 | CLDN-18 | *Scophthalmus_maximus* |
| 55 | XP_003444253.1 | CLDN18 | *Oreochromis_niloticus* |
| 56 | XP_030228284.1 | CLDN18 | *Gadus_morhua* |
| 57 | XP_018586618.1 | CLDN-18-like | *Scleropages_formosus* |
| 58 | XP_035278220.1 | CLDN18 | *Anguilla_anguilla* |
| 59 | XP_012681094.1 | CLDN18 | *Clupea_harengus* |
| 60 | XP_014045392.1 | CLDN18 | *Salmo_salar* |
| 61 | XP_002660862.5 | CLDN18 | *Danio_rerio* |
| 62 | XP_010891673.1 | CLDN18 | *Esox_lucius* |
| 63 | XP_026882749.2 | CLDN18 | *Electrophorus_electricus* |
| 64 | NP_001316217.1 | claudin_18 | *Ictalurus_punctatus* |
| 65 | XP_029930832.1 | CLDN18-like | *Myripristis_murdjan* |
| 66 | XP_030220814.1 | CLDN18-like | *Gadus_morhua* |
| 67 | XP_040027686.1 | CLDN18 | *Gasterosteus_aculeatus_aculeatus* |
| 68 | XP_003438075.1 | CLDN18-like | *Oreochromis_niloticus* |
| 69 | XP_018532695.1 | CLDN-18 | *Lates_calcarifer* |
| 70 | XP_011614961.1 | CLDN18 | *Takifugu_rubripes* |
